# Multiple macroecological laws do characterize various aspects of microbiota dynamics

**DOI:** 10.1101/2021.08.09.455744

**Authors:** Konstantine Tchourine, Martin Carballo-Pacheco, Dennis Vitkup

## Abstract

In this letter we address the potential confusion related to our recent demonstration that multiple macroecological laws describe short- and long-term dynamics of microbial communities. Specifically, we clarify that these laws, similarly to many other relationships observed in nature, are characterized not just by the existence of scaling, but also by certain characteristic values of the scaling exponents. By performing proper statistical analysis, we demonstrate that the relationships sensitive to temporal bacterial dynamics are not reproduced in the shuffled data. We also discuss that there is no clear evidence in the data that macroecological relationships in microbiota are primarily driven by external or environmental factors. Proper statistical analyses of the data suggest that the dynamics of gut microbiota, even on a constant diet, contains rich temporal structure. Therefore, it is likely that complex and non-linear internal dynamics may be primarily responsible for the observed macroecological laws in microbiota and other ecological communities.

---

We are surprised by the letter by Wang and Liu^1^ related to our recent study demonstrating that short- and long-term microbiota dynamics can be characterized by multiple macroecological relationships^2^. The discovery of these laws in microbiota represents an important advance in the understanding of microbial ecology. Our results suggest that similar underlying principles may govern the dynamics of both micro- and macroecological communities. Moreover, we demonstrate that the macroecological relationships are predictive and can be used to identify specific bacterial strains with abnormal or perturbed dynamics. Wang and Liu appear to be confused about the nature of our analysis and about the origin of the macroecological laws in general. Furthermore, there are several unfortunate technical errors in their analysis as well as some expected mathematical observations. To prevent misunderstanding in the field, we address in this letter both the conceptual and the technical issues raised by Wang and Liu.

On the conceptual side, the macroecological laws describe statistical relationships between various features of microbiota dynamics. By analyzing shuffled time series of bacterial dynamics, Wang and Liu suggest that: 1) microbiota dynamics and macroecological relationships must be primarily externally driven, and 2) macroecological laws may not reflect the real dynamics of microbiota. We disagree with these claims, and we have difficulty understanding the logic of their analysis. First, it seems self-evident that some macroecological relationships may be relatively insensitive to the ordering of bacterial abundance trajectories (see below), while still accurately describing species’ *dynamics*, i.e., “a pattern or process of change, growth, or activity”. Second, in ecology in general, and in our analysis in particular, these relationships characterize population dynamics but make no particular claims about the origin of the dynamics, such as whether they are stochastic or deterministic, driven by external perturbations, internal perturbations, or some combination of both. Under all of these scenarios, the macroecological laws faithfully describe bacterial dynamics. Third, the conclusion that because similar functional forms of distributions are often observed in shuffled data, microbiota dynamics must be primarily externally driven is not correct. More than 40 years ago, the pioneering work of Robert May^3-5^ demonstrated that simple non-linear ecological systems, often driven by their complex internal dynamics, can result in chaotic and unpredictable temporal behavior. Therefore, complex dynamics of ecological systems usually cannot be predicted by simple autoregressive temporal models. These insights have been confirmed by many researchers and now represent classic and well-established results in the field of ecology^6^.

On the technical side, we would like to emphasize that shuffled trajectories *do not represent random bacterial dynamics* in the context of studying macroecological laws. Many of the relevant features present in the real data, and sometimes all the relevant features, will be necessarily preserved in the shuffled trajectories. By performing proper statistical analysis, we demonstrate below that the relationships crucially dependent on temporal dynamics are in fact not reproduced in the shuffled data. The functional form, but not the characteristic parameters of the relationships that only weakly depend on temporal dynamics, appear similar, as they predictably should, although shuffled distributions are again statistically different from the distributions in the real data. Finally, the laws that do not depend on day-to-day dynamics are reproduced exactly, again as they mathematically should be. Notably, in our original paper, we use a null model based on random sampling of sequencing data to show (Figure 1D below) that the observed temporal fluctuations, both in terms of their magnitude and their form, are not due to sampling noise in the data.

**Figure 1.**
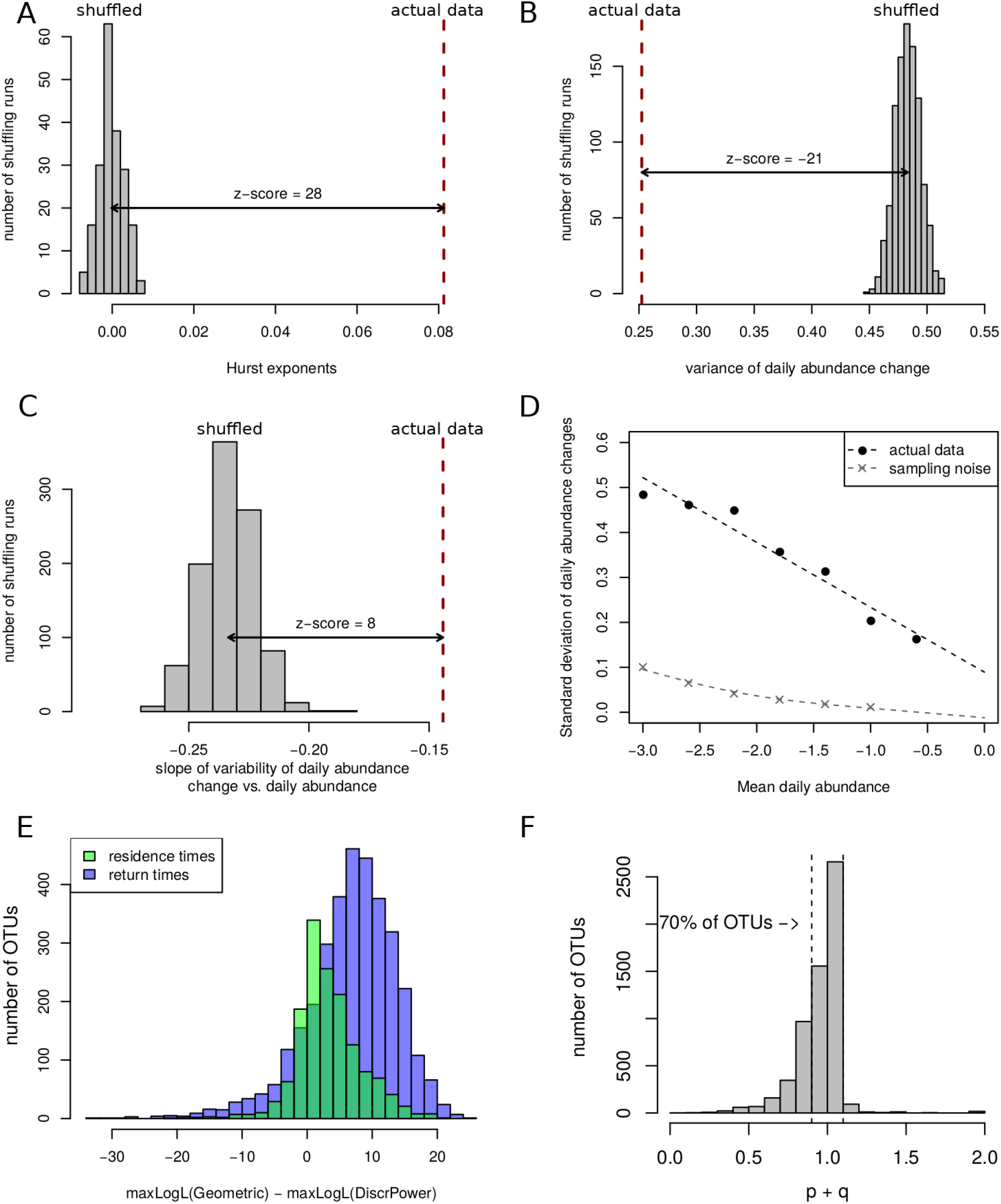
Analysis of the time-shuffling data transformation. **A**. Distribution of Hurst exponents across time-shuffling runs. **B**. Distribution of variances of daily abudnance changes across time-shuffling runs. **C**. Distribution of slopes of the linear relationship between standard deviation (s.d.) of daily abundance change and mean daily abundance. **A-C**. The distributions of the scaling law statistics were calculated across 200 (**A**) and 1000 (**B-C**) random temporal shufflings of the data. Red vertical dashed lines denote the corresponding measurments in the real biological data. Z-scores indicate the number of standard deviations between the means of the shuffled distributions and the values in the real data; in all cases, the values in the real data are substantially and signfiicnatly different from the time-shuffled distribution (***p*** < **10**^**−5**^). **D**. Relationship between the s.d. of daily abundance changes and mean daily abundance for the original data (black circles) and for the random sampling null model (grey crosses). Black dashed line shows best line of fit to the original data. **E**. The difference between the maximum log-likelihood (maxLogL) of the geometric distribution fits to the residence (green) or return (blue) data for each OTU and the maximum log-likelihoods of the power law distribution fits to the data for the same OTU; exactly one parameter was fit for each model for each OTU (see Materials and Methods). In both cases, the geometric distribution is significantly more likely to describe the behavior for the majority of OTUs (paired t-test, p<**10**^**−10**^). **F**. Distribution of the sum of extinction probability ***p*** and reemergence probability ***q***, fitted to each OTU individually, assuming a geometric distribution of residence and return times evaluated at its maximum likelihood. Percent of OTUs with **0. 9** < ***p*** + ***q*** < **1. 1** is shown in the figure. For illustrative purposes, only the data from the Human A dataset is shown in all panels, but similar results are obtained for all human and mouse datasets we analyzed in the paper.

Shuffling the gut microbial time series results in several trivial observations consistent with the original data structure. For example, in the original work we showed that the variance of species (OTU) abundance is related to the mean through a power law, a relationship often referred to in ecology as Taylor’s law^7-10^. Taylor’s law allowed us to identify outliers in bacterial dynamics, including OTUs related to environmental and health-related perturbations. Wang and Liu show that shuffling the temporal data does not affect Taylor’s law. Obviously, the average and the variance of a set of numbers, in this case bacterial abundances and their variances over time, are not affected by reordering of the numbers. We do not understand what is surprising here.

We demonstrated that long-term microbiota dynamics can be characterized by Hurst diffusion in log OTU abundance space. Specifically, the mean-squared displacement of log abundance ratios is proportional to the diffusion time: ⟨*δ*^2^(Δ*t*)⟩ ∝Δ*t*^2*H*^. We showed that for entire microbiota and for many individual OTUs, *H* ≈ 0.1 − 0.2. We performed shuffling of microbiota trajectories and found that the Hurst exponents observed in the real data are significantly higher than for the time-shuffled data (Figure 1A; *Z* > 19 for human datasets, and *Z* > 5 for mouse datasets; *p* < 10^−5^ for all datasets), where the Hurst exponent is *H* ≈ 0. This is expected, as the shuffling destroys temporal correlation in the data, thereby effectively transforming a sub-diffusive process into a mean-reverting process. However, the claim by Wang and Liu that this is an example of the same law in the shuffled data is perplexing, because scientific relationships are not simply characterized by certain functional forms or scaling, but also by specific values of key parameters of corresponding relationships.

We also showed that daily abundance changes follow a Laplace distribution, a relationship previously observed in many other ecosystems. Wang and Liu find that the shuffled data also follow a Laplace distribution, but with larger exponents and therefore larger distribution variances. The observed difference in exponents and variances is not surprising, because for two consecutive measurements in the real data, OTU abundances are significantly more similar than for two randomly selected time points in the shuffled trajectories. This again demonstrates clear temporal structure in the data. Furthermore, we found that the variance of daily abundance changes in the real data is substantially and significantly lower (*Z* < −9; *p* < 10^−5^ for all datasets) than for the shuffled data in all human and mouse datasets (Figure 1B).

A similar argument applies in the case of the linear relationship between the variability of daily abundance changes *σ*_*μ*_ and mean daily abundance *x*_*m*_. This relationship demonstrates that abundant bacterial species display predictably lower relative fluctuations compared to species with lower abundances. The general similarity of the relationship in shuffled data is again expected, as the average abundance for a given OTU tends to remain similar during the entire time series, i.e., high/low abundant species tend to remain high/low abundant across temporal trajectories. Therefore, temporal shuffling results in a similar functional relationship. By analyzing shuffled trajectories, we again found a statistically significant difference in the slope parameter compared to the real data in all considered datasets (Figure 1C; *Z* > 5; *p* < 10^−5^). Moreover, as we demonstrated in the original paper, the fluctuations described by this relationship are not due to sampling noise (Figure 1D).

We also demonstrated that the distributions of bacterial residence and return times across species follow a power law with specific parameters, and Wang and Liu find that these distributions do not substantially change following temporal shuffling. This observation is yet again a mathematical necessity. For individual OTUs, residence and return times are consistent with the geometric distribution (Figure 1E). Exponentially (geometrically) distributed extinction and lifespan times have been observed for many individual species in ecology^11-13^. Furthermore, the probability of extinction (*p*) and reemergence (*q*) at each step for individual bacterial species are related such that *p* + *q* ≈ 1 (see Figure 1F). This corresponds to a process in which the absence and presence of each species is controlled by a binary random variable, with probability *p* of absence at each time step, and probability 1 − *p* = *q* of presence at each time step. This again does not imply that an OTU’s presence and absence are necessarily determined by external stochastic noise, as chaotic dynamics can often result in similar behavior. From such a process, it follows that *p* will be approximately equal to the fraction of time points at which the species is absent, and *q* will approximate the fraction of time points for which it is present. Shuffling temporal data with an approximately constant probability of presence and absence at each time step will then necessarily result in a similar distribution of the residence and return times for individual OTUs as in the original data. The distributions of these residence and return times across species will then also be identical, exactly as Wang and Liu observed.

Finally, Wang and Liu computed the noise color profile and concluded that most of the temporal variance is white noise. The results of this analysis were in part behind their claim of “very weak temporal structure” in the data. However, it appears that Wang and Liu decomposed the noise for all OTUs, whereas in the paper, following the convention in the field, we analyzed the dynamics of bacterial abundance using only highly abundant bacteria. In fact, our group recently demonstrated, using careful experimental and computational analysis^14^, that the variance of OTUs with a relative abundance below ∼10^−3^ is often dominated by technical noise, while the variance of OTUs above this threshold is dominated by biologically-relevant temporal and spatial variation. Thus, the white noise observed by Wang and Liu simply reflects technical noise in the lowly-abundant OTUs, which represent over 97% of all OTUs in each dataset. Thus, their conclusions are not accurate, and the results are again expected. We reanalyzed the data considering only the highly abundant OTUs (analyzed in the original paper) and found that most noise profiles are actually dominated by pink (1/f) noise, which often indicates long-term memory in temporal trajectories^15^, whereas shuffled data predictably results in white noise (Figure 2); mouse time series were significantly shorter than human datasets, which resulted in more white noise in their spectra. We note that results similar to ours are also obtained by Faust et al.^16^, whom Wang and Liu in fact cite. This analysis reinforces the conclusion that microbiota dynamics often contains a rich and relevant temporal structure.

**Figure 2.**
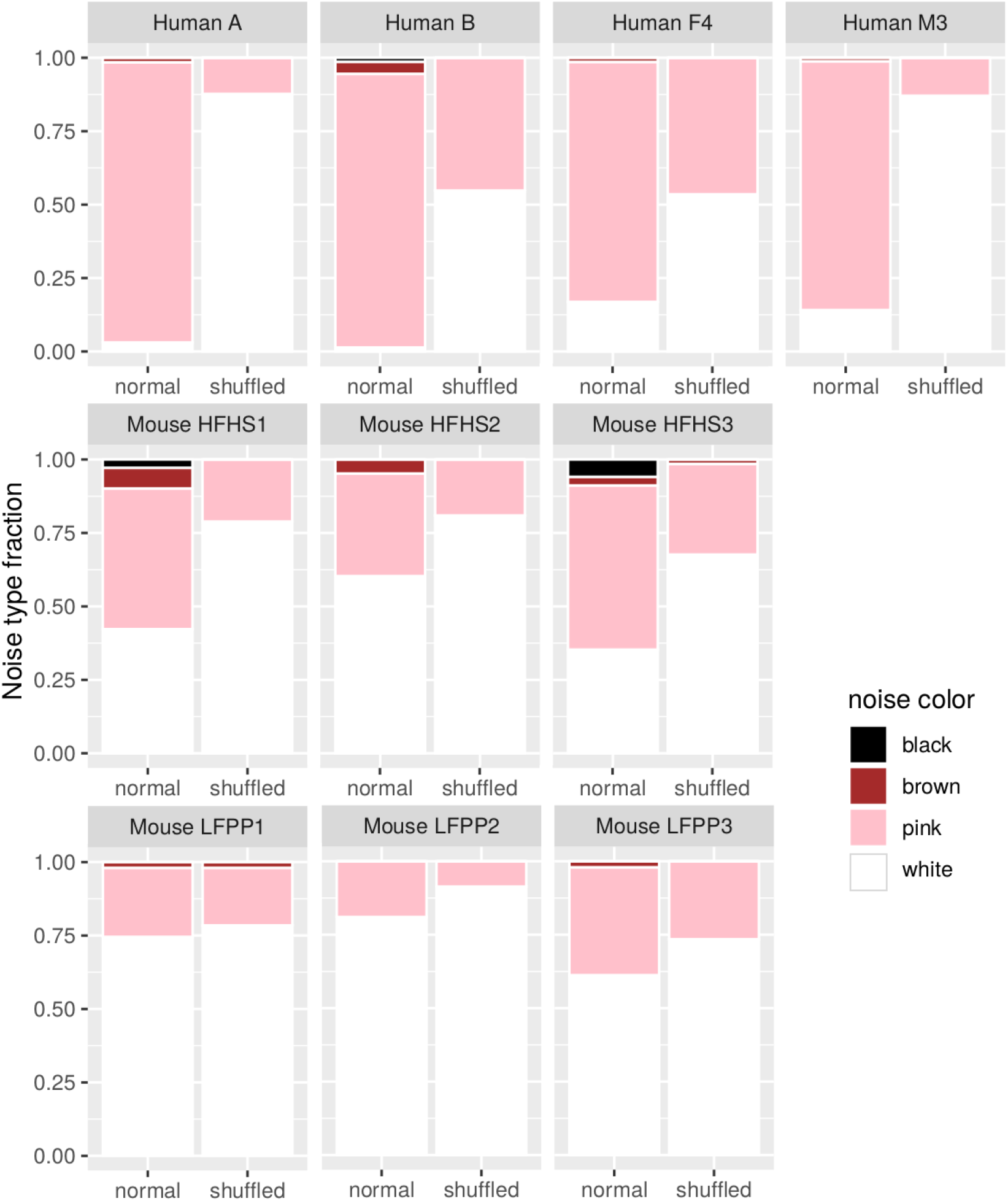
Noise decomposition profiles for human and mouse datasets. Consistent with the OTUs used for the analysis of abundance dynamics in Ji *et al*., 2020, the spectral noise decomposition was performed only for the OTUs with average abundance over **10**^**−3**^. The approach for noise decomposition and definitions of noise colors from Faust *et al*. ^16^ were used (see Materials and Methods).

In conclusion, understanding the nature of macroecological relationships in microbiota and other ecological systems is an important and exciting research direction. Therefore, it is essential to start exploring these laws without confusion about the presented results and about existing ecological literature. We think an important next step will be to develop models capable of robustly fitting experimental macroecological relationships, including scaling exponents observed in real data rather than just scaling itself, and models capable of fitting all macroecological laws simultaneously rather than only a fraction or some elements of the observed laws. Moreover, models that are based on known and observable features of microbiota dynamics will be more useful than models invoking additional sources of external or internal noise to fit the data. We note in this respect that, as we have demonstrated previously^2^, macroecological laws are also observed in mice on constant diets. Therefore, although diets do affect microbiota dynamics, it is unlikely that food or environmental variability is primarily responsible for the observed macroecological patterns, at least in general. We believe that further experimental and modeling studies that take into account the aforementioned observations will provide valuable mechanistic insights into ecological dynamics of microbiota and beyond.

## Materials and Methods

### Scaling law calculations

Human^17,18^ and mouse^19^ data sets were processed identically as in Ji *et al*., (2020)^2^. In order to control for technical factors, a cut-off of mean relative abundance of 10^−3^ and a requirement to be present in half of the time series was used to filter out low-abundance OTUs for each dataset. Each scaling law was calculated identically to Ji *et al*., (2020)^2^.

### Maximum likelihood fits to the distributions of residence and return times

We calculated the log-likelihood of the geometric distribution for each OTU using its standard formula. Specifically, the log-likelihood of the geometric distribution is given by 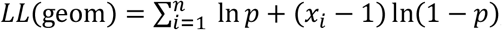, where *p* is a parameter representing the probability of extinction (reemergence), and *x*_*i*_ represents the vector of measured presence (absence) times. Maximum likelihood values were calculated by analytically solving for the optimal parameter *p*. Discrete power law fits were calculated by maximizing the log-likelihood function 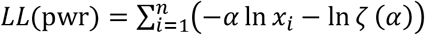, where *α* is the power law scaling parameter, *x*_*i*_ is the vector of measured presence (absence) times, and *ζ*(*α*) is the Riemann zeta function evaluated at *α*^20^. We maximized *LL*(pwr) by performing a grid search on the parameter *α* ranging from 0 to 10 at intervals of 0.01.

### Noise color calculations

To calculate the noise color for each OTU in each dataset, we followed the exact procedure given in Faust *et al*. (2018)^16^. First, missing time points for each trajectory were interpolated by Stineman interpolation using the R package stinepack^21^. Frequency and spectral density were then calculated with detrending enabled. Spectral density was then fit with a spline in log-log scale as a function of frequency, setting the degree of freedom to the maximum of [2, log10(length of the time series)], using the R function smooth.spline. The minimum of the first derivative of the resulting fit was used as a measure of the relationship between frequency and spectral density of the time trajectory. This value was used to classify each OTU’s color of the noise as either black (below −2.25), brown (between −2.25 and −1.75), pink (between −1.75 and −0.5), or white (over −0.5), following the scheme given in *Faust et al*. (2018)^16^. Only the OTUs that passed the abundance and prevalence cutoffs from Ji *et al*. (2020)^2^ were used for this calculation.

## References

1 Wang, X.-W. & Liu, Y.-Y. Characterizing scaling laws in gut microbial dynamics from time series data: caution is warranted. bioRxiv, 2021.2001.2011.426045, doi:10.1101/2021.01.11.426045 (2021).

2 Ji, B. W., Sheth, R. U., Dixit, P. D., Tchourine, K. & Vitkup, D. Macroecological dynamics of gut microbiota. Nature Microbiology 5, 768–775, doi:10.1038/s41564-020-0685-1 (2020).

3 May, R. M. Simple mathematical models with very complicated dynamics. Nature 261, 459–467, doi:10.1038/261459a0 (1976).

4 May, R. M. Biological populations with nonoverlapping generations: stable points, stable cycles, and chaos. Science 186, 645–647, doi:10.1126/science.186.4164.645 (1974).

5 May, R. M. Biological populations obeying difference equations: Stable points, stable cycles, and chaos. Journal of Theoretical Biology 51, 511–524, doi:10.1016/0022-5193(75)90078-8 (1975).

6 May, R.M.R.M. & McLean, A. R. Theoretical ecology: principles and applications. 98–132 (2007).

7 Taylor, L. R. Aggregation, Variance and the Mean. Nature 189, 732–735, doi:10.1038/189732a0 (1961).

8 Taylor, L. R. & Woiwod, I.P. Temporal Stability as a Density-Dependent Species Characteristic. The Journal of Animal Ecology 49, 209–224, doi:10.2307/4285 (1980).

9 Taylor, L. R., Woiwod, I. P. & Perry, J.N. The Density-Dependence of Spatial Behaviour and the Rarity of Randomness. The Journal of Animal Ecology 47, 383–406, doi:10.2307/3790 (1978).

10 Eisler, Z., Bartos, I. & Kertesz, J. Fluctuation scaling in complex systems: Taylor’s law and beyond. Adv. Phys. 57, 89–142, doi:10.1080/00018730801893043 (2008).

11 Pigolotti, S., Flammini, A., Marsili, M. & Maritan, A. Species lifetime distribution for simple models of ecologies. Proceedings of the National Academy of Sciences of the United States of America 102, 15747–15751, doi:10.1073/pnas.0502648102 (2005).

12 Shimada, T., Yukawa, S. & Ito, N. Life-span of families in fossil data forms q-exponential distribution. International Journal of Modern Physics C 14, 1267–1271, doi:10.1142/S0129183103005406 (2003).

13 Solé, R.V. & Bascompte, J. Are critical phenomena relevant to large-scale evolution? Proceedings of the Royal Society of London. Series B: Biological Sciences 263, 161–168, doi:10.1098/rspb.1996.0026 (1996).

14 Ji, B. W. et al. Quantifying spatiotemporal variability and noise in absolute microbiota abundances using replicate sampling. Nature Methods 16, 731–736, doi:10.1038/s41592-019-0467-y (2019).

15 Graves, T., Gramacy, R., Watkins, N. & Franzke, C. A Brief History of Long Memory: Hurst, Mandelbrot and the Road to ARFIMA, 1951–1980. Entropy 19, 437, doi:10.3390/e19090437 (2017).

16 Faust, K. et al. Signatures of ecological processes in microbial community time series. Microbiome 6, 120, doi:10.1186/s40168-018-0496-2 (2018).

17 David, L. A. et al. Host lifestyle affects human microbiota on daily timescales. Genome Biology 15, 1–15, doi:10.1186/gb-2014-15-7-r89 (2015).

18 Caporaso, J. G. et al. Moving pictures of the human microbiome. Genome Biology 12, 1–8, doi:10.1186/gb-2011-12-5-r50 (2011).

19 Carmody, R. N. et al. Diet dominates host genotype in shaping the murine gut microbiota. Cell Host and Microbe 17, 72–84, doi:10.1016/j.chom.2014.11.010 (2015).

20 Clauset, A., Shalizi, C. R. & Newman, M. E. J. Power-Law Distributions in Empirical Data. SIAM Review 51, 661–703, doi:10.1137/070710111 (2009).

21 Stineman, R. A consistently well-behaved method of interpolation. Creative Computing 6, 54–57 (1980).

